# NeuroNLP: a natural language portal for aggregated fruit fly brain data

**DOI:** 10.1101/092429

**Authors:** Nikul H. Ukani, Adam Tomkins, Chung-Heng Yeh, Wesley Bruning, Allison L. Fenichel, Yiyin Zhou, Yu-Chi Huang, Dorian Florescu, Carlos Luna Ortiz, Paul Richmond, Chung-Chuan Lo, Daniel Coca, Ann-Shyn Chiang, Aurel A. Lazar

**Affiliations:** Department of Electrical Engineering, Columbia University, New York, NY 10027, USA; Department of Automatic Control & Systems Engineering, The University of Sheffield, Sheffield, S1 3JD, UK; Department of Computer Science, Columbia University, New York, NY 10027, USA; Data Science Institute, Columbia University, New York, NY 10027, USA; Brain Research Center, National Tsing Hua University, Hsinchu 30013, Taiwan; Institute of Systems Neuroscience, National Tsing Hua University, Hsinchu 30013, Taiwan; Department of Life Science, National Tsing Hua University, Hsinchu 30013, Taiwan; Department of Computer Science, The University of Sheffield, Sheffield, S1 4DP, UK; Genomics Research Center, Academia Sinica, Nankang, Taipei 11529, Taiwan; Institute of Physics, Academia Sinica, Nankang, Taipei 11529, Taiwan; Department of Biomedical Science and Environmental Biology, Kaohsiung Medical University, Kaohsiung 80708, Taiwan; Kavli Institute for Brain and Mind, University of California, San Diego, La Jolla, California 92093, USA

## Abstract

NeuroNLP, is a key application on the Fruit Fly Brain Observatory platform (FFBO, http://fruitflybrain.org), that provides a modern web-based portal for navigating fruit fly brain circuit data. Increases in the availability and scale of fruit fly connectome data, demand new, scalable and accessible methods to facilitate investigation into the functions of the latest complex circuits being uncovered. NeuroNLP enables in-depth exploration and investigation of the structure of brain circuits, using intuitive natural language queries that are capable of revealing the latent structure and information, obscured due to expansive yet independent data sources. NeuroNLP is built on top of a database system call NeuroArch that codifies knowledge about the fruit fly brain circuits, spanning multiple sources. Users can probe biological circuits in the NeuroArch database with plain English queries, such as “show glutamatergic local neurons in the left antennal lobe” and “show neurons with dendrites in the left mushroom body and axons in the fan-shaped body”. This simple yet powerful interface replaces the usual, cumbersome checkboxes and dropdown menus prevalent in today’s neurobiological databases. Equipped with powerful 3D visualization, NeuroNLP standardizes tools and methods for graphical rendering, representation, and manipulation of brain circuits, while integrating with existing databases such as the FlyCircuit. The userfriendly graphical user interface complements the natural language queries with additional controls for exploring the connectivity of neurons and neural circuits. Designed with an open-source, modular structure, it is highly scalable/flexible/extensible to additional databases or to switch between databases and supports the creation of additional parsers for other languages. By supporting access through a web browser from any modern laptop or smartphone, NeuroNLP significantly increases the accessibility of fruit fly brain data and improves the impact of the data in both scientific and educational exploration.

## Additional Details

### Workflow of NeuroNLP

The architecture of the FFBO [1] is sketched in Figure 1. NeuroNLP utilizes two main modules in the architecture, namely, the natural language processing (NLP) module and the NeuroArch Database. NeuroArch [2] is a novel, graph-based database that supports the integration of fruit fly brain data from multiple sources. The NLP module was developed to parse English queries into NeuroArch API calls. NeuroArch currently stores fly connectome data from two publicly available sources: the FlyCircuit database [3] and the Janelia Fly Medulla data [4], and is open to support additional data. The former hosts meso-scale connectome data on the whole-brain level and the latter contains detailed, micro-scale data about exact chemical. synaptic information between neurons in a limited region of the Medulla neuropil. Combining data from multiple sources into a single database, with a common data model, NeuroNLP facilitates the access and retrieval of data from various sources simultaneously using the same queries.

**Figure 1:**
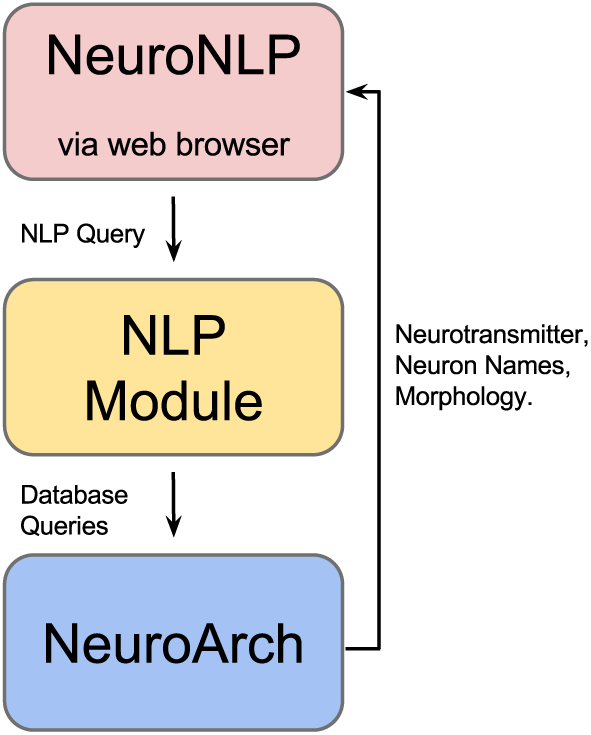
Basic building blocks and workflow of NeuroNLP.

### Supported Query Types

NeuroNLP currently supports queries to show, add, filter and remove neurons based on 1) the neuropil that neurons belongs to, 2) the type of neuron (either projection neuron or local neuron), 3) dendritic/axonal arborization, 4) neurotransmitter type and 5) postsynaptic and presynaptic neurons in relation to a set of neurons.

**Figure 2:**
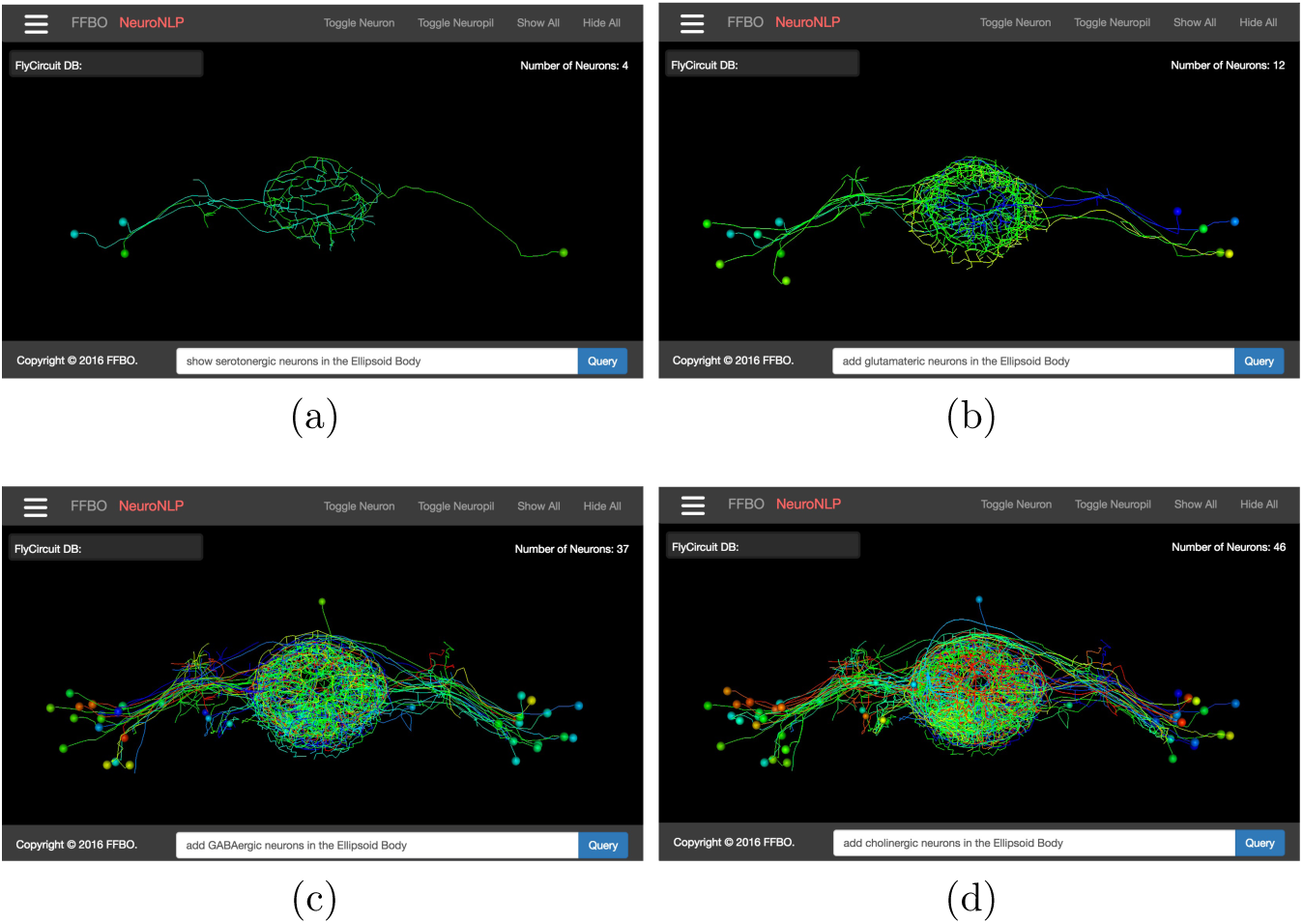
A sequence of queries to reveal ring neurons with different types of neurotransmitter in the EB. (a) Serotonergic (b) Glutamatergic (c) GABAergic (d) Cholinergic.

### Examples

Ring neurons in the Ellipsoid Body (EB) have been characterized by their prominent ring structure but their functional role has not been understood. By using a sequence of queries: “show serotonergic neurons in EB, add glutamatergic/GABAergic/cholinergic neurons in EB”), we can easily obtain a neurotransmitter profile of the ring neurons in the EB. The result suggests that a diverse population of ring neurons exists in the EB that may differentially contribute to circuit function. Such a quick search can immediately inspire/lead to new questions.

It is known that the Lobula Plate Tangential Cells (LPTCs) are an essential element in the fly optomotor pathway. Most of the literature shows that LPTCs have extensive dendritic arborizations in the lobula plate (LOP) and terminate in the caudal ventrolateral protocerebrum (CVLP), with their cell body in the rind of LOP. Using a few quick query and GUI operations, we found 16 neurons that have large arborization in the LOP. Out of the 16, 7 were not previously described, as they either have terminals on both sides of CVLP or have a cell body located at the center of the brain. These findings can lead to additional investigations of the circuit structure of the optomotor pathway as well as its function in motion detection.

**Figure 3:**
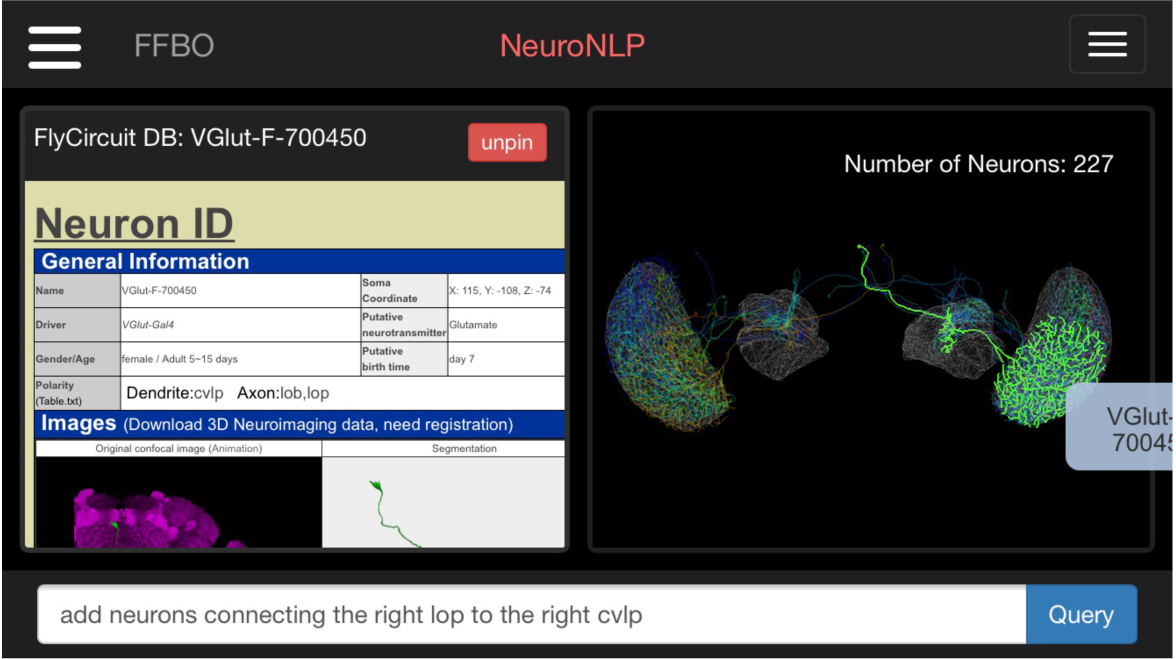
16 LPTCs extracted from the database using NeuroNLP on an iPhone 6s. One of the neuron is highlighted, and its information page on the FlyCircuit database is shown on the left.

Streamlining the way we simultaneously explore and interrogate distal data sources can open up new avenues of research, and enrich neuroscience education.

